# Deletion of *FUNDC2* and *CMC4* on chromosome Xq28 is sufficient to cause hypergonadotropic hypogonadism in men

**DOI:** 10.1101/2020.03.23.004424

**Authors:** Xinxian Deng, He Fang, Asha Pathak, Angela M. Zou, Whitney Neufeld-Kaiser, Emily A. Malouf, R. Alan Failor, Fuki M. Hisama, Yajuan J. Liu

## Abstract

**Background:** Hypergonadotropic hypogonadism (HH) is characterized by low sex steroid levels and secondarily elevated gonadotropin levels with either congenital or acquired etiology. Genetic factors leading to HH have yet to be fully elucidated.

**Methods:** Here, we report on genome and transcriptome data analyses from a male patient with HH and history of growth delay who has an inherited deletion of chromosome Xq28. Furthermore, expression analyses were done for this patient and his unaffected family members and compared to normal controls to identify dysregulated genes due to this deletion.

**Results:** Our patient’s Xq28 deletion is 44,806bp and contains only two genes *FUNDC2* and *CMC4*. Expression of both *FUNDC2* and *CMC4* are completely abolished in the patient. Gene ontology analyses of differentially expressed genes in the patient in comparison to controls show that significantly up-regulated genes in the patient are enriched in Sertoli cell barrier regulation, apoptosis, inflammatory response and gonadotropin-releasing regulation. Indeed, our patient has an elevated FSH level, which regulates Sertoli cell proliferation and spermatogenesis. In his mother and sister, who are heterozygous for this deletion, X-chromosome inactivation is skewed towards the deleted X, suggesting a mechanism to avoid FSH dysregulation.

**Conclusion:** Compared to the previously reported men with variable sized Xq28 deletions, our study suggests that loss of function of *FUNDC2* and/or *CMC4* results in dysregulation of apoptosis, inflammation and FSH, and is sufficient to cause Xq28-associated HH.

## 1 INTRODUCTION

Hypergonadotropic hypogonadism (HH; also termed primary hypogonadism) in men is defined by reduced production of either sperm, testosterone, or both, accompanied by elevated levels of the pituitary gonadotropins luteinizing hormone (LH) and follicle stimulating hormone (FSH) (Basaria, 2014; Viswanathan & Eugster, 2011). LH is required for Leydig cell (LC) hyperplasia and testosterone release in males. FSH regulates Sertoli cell (SC) proliferation before and at puberty, and participates in the regulation of spermatogenesis. HH can be primary, caused by constitutional chromosomal abnormalities or smaller genetic variants, or secondary, due to autoimmunity or exposure to chemotherapy/radiation.

The most common constitutional cause of HH in both men and women is X chromosome abnormalities, including Turner syndrome (45,X), 47,XXY, 47,XXX and Xq deletion (Viswanathan & Eugster, 2011). This strongly suggests that correct dosage of some X-linked genes is critical in normal gonad development. Indeed, the mammalian X chromosome was specifically enriched in genes involved in reproduction during the evolution of the sex chromosomes from a pair of ordinary autosomes (Deng et al., 2014). Which X-linked genes are involved in gonad development has yet to be fully elucidated. Genetic evaluation of patients with HH and X-chromosome abnormalities, especially chromosomal microdeletions spanning a few genes, may help identify candidate genes and provide new insights of the etiology of HH.

Here we report a 35-year-old man with HH, short stature, and bilateral cataracts who was identified have a 44.8 kb deletion of chromosome Xq28 encompassing the entirety of *FUNDC2 (FUN14 Domain Containing* 2), which encodes a mitochondrial membrane protein, and all but the shared exon 1 of *CMC4* (*C-X9-C motif containing 4*) and *MTCP1* (*mature T cell proliferation 1*). *MTCP1* encodes p13^MTCP1^, which is absent in normal tissue and is detected only in a rare T-cell prolymphocytic leukemia with t(X;14) translocations (Madani et al., 1996). Thus *MTCP1* is probably dispensable for gonad development. *CMC4* shares a common promoter and 5’ UTR (exon 1) with *MTCP1* but has a distinct set of coding exons. *CMC4* is expressed in many tissues and encodes p8^MTCP1NB^, a mitochondrial membrane protein of unknown function with highest level in fetal testis as shown in HIPED (the database of protein abundance in human tissues) (Fagerberg et al., 2014; Fishilevich et al., 2016; Madani et al., 1995).

Mouse knockout studies have shown that loss of function of *FUNDC2* induces platelet apoptosis under hypoxic stress and impairs platelet activation and aggregation (Ma, Zhang, Zhu, Liu, & Chen, 2019; Ma, Zhu et al., 2019). To investigate the roles of *CMC4* and *FUNDC2* in gonad development and the pathways leading to HH in this patient, expression analyses including RNA-seq were done for this patient and his unaffected family members, including his heterozygous mother. In comparison to expression data from control males and females, we found that significantly up-regulated, but not down-regulated, genes in the patient are enriched in SC regulation, apoptosis and inflammatory response, and gonadotropin-releasing regulation. Dysregulation of SCs, and the gonadotropin-releasing pathway in the patient is consistent with the clinical diagnosis of HH. Interestingly, it has been proposed that increased apoptosis affecting SCs and LCs is a main mechanism leading to testis dysfunction in men with 47,XXY (D’Aurora et al., 2015; D’Aurora et al., 2017). In our patient’s mother and sister, who are heterozygous for the deletion, X-chromosome inactivation (XCI) is highly skewed towards the deleted X chromosome, suggesting a mechanism in females to avoid FSH dysregulation and defects in ovarian follicular maturation in early development. Xq28 deletions have previously been reported in only 8 men with syndromic forms of HH (Lavin et al., 2016; Miskinyte et al., 2011). Compared to these studies, our study suggests that loss of function of *FUNDC2* and *CMC4* leads to increased apoptosis and inflammation and damaged SC functionality, which probably causes Xq28-associated HH.

## 2 MATERIALS AND METHODS

### 2.1 Protocol approvals and patient consents

After written, informed consent was obtained from the patient and each of his unaffected family members, including his mother, sister, brother, and both maternal uncles (Figure S1), they were enrolled in this study under a protocol approved by the institutional review board of University of Washington. Peripheral blood samples or saliva sample were obtained for DNA, and RNA studies.

### 2.2 Cytogenetics and Chromosomal SNP Microarray Analysis

G-banded chromosome analysis and karyotyping were done by standard methods. Interphase fluorescence in situ hybridization (FISH) analysis was performed on peripheral blood cells from the patient using probes for chromosomes X (DXZ1) and Y (DYZ3). Chromosomal SNP (single nucleotide polymorphism) microarray analysis of genomic DNA prepared from peripheral blood (the patient, his mother, his brother, and one maternal uncle) or from saliva (his sister, one maternal uncle) was performed using the Illumina Infinium CytoSNP-850K BeadChip v1.1. Microarray data was visualized and analyzed using Illumina BlueFuse Multi v4.4.

### 2.3 PCR and Sanger sequencing

To map the breakpoints of the patient’s deletion, DNA was isolated from peripheral blood and amplified by PCR (HotStart II High Fidelity PCR Master Mix, Thermo Fisher Scientific), followed by Sanger sequencing (Eurofins Genomics). The PCR primers used were: TCAAAATAGACCCCATTACCAAA (forward primer); GGGGACAGCCCTTTAAGACG (reverse primer).

### 2.4 mRNA-seq

RNA-seq experiments were conducted on samples from the patient and 3 unaffected family members (mother, one maternal uncle, and brother) at QIAGEN Genomic Services. The library preparation was done using TruSeq^®^ Stranded mRNA Sample preparation kit (Illumina). In brief, the starting material (100ng) of total RNA isolated from peripheral blood was mRNA enriched using the oligodT bead system. The isolated mRNA was subsequently fragmented using enzymatic fragmentation. Then first strand synthesis and second strand synthesis were performed and the double stranded cDNA was purified (AMPure XP, Beckman Coulter). The cDNA was end repaired, 3’ adenylated and Illumina sequencing adaptors ligated onto the fragments ends, followed by PCR amplification and purification. The libraries size distribution was validated and quality inspected on a Bioanalyzer tapeStation (Agilent Technologies). High quality libraries are pooled based in equimolar concentrations based on the Bioanalyzer Smear Analysis tool (Agilent Technologies). The library pool(s) were quantified using qPCR and optimal concentration of the library pool used to generate the clusters on the surface of a flowcell before 75nt paired-end sequencing on a NextSeq500) instrument (2×75 cycles) according to the manufacturer instructions (Illumina).

Reads were aligned to the human annotation reference genome GRCh38 using STAR 2.5.2b. BAM files of mapped reads were visualized by the Integrative Genomics Viewer (IGV). ~26million (M) uniquely mapped reads (mappability: >84%) were obtained for the patient and mother, respectively. However, only 19 M (mappability: 68%) and 8 M (mappability: 61%) uniquely mapped reads were obtained for uncle and brother, respectively. The low mappability suggests that the RNA samples from uncle and brother was degraded (i.e. fewer reads will be mRNA or lncRNA specific and more material will be degraded rRNA and mtRNA). Indeed, RNA integrity number (RIN) is very low (RIN <3) for the RNA samples from uncle and brother while RIN is >9 for the RNA samples from the patient and mother. Thus only RNA-seq data from the patient and mother were used for the differential expression (DE) analysis in comparison to public mRNA-seq data sets of three males (SRR9666161 (age 33), SRR9666180 (age 32), SRR9666239 (age 40)) and three females (SRR9666175 (age 28), SRR9666197 (age 43), SRR9666240 (age 28)) with age matched to the patient (RNA-seq whole blood of Dutch 500FG cohort, GSE134080). These control data sets have a similar sequence depth and quality with 13-17 M uniquely mapped reads (mappability: >87%). Raw counts of the Genecode genes from STAR 2.5.2b alignment are present in **Dataset 1**, which is used for expression cutoffs for the DE analysis.

The DE analyses were done with weighted trimmed mean of M-values (TMM) normalization method and Genecode gene annotation in the EdgeR statistical software package (Bioconductor, http://www.bioconductor.org/) to investigates the relative change in gene expression (i.e. normalized counts) between different samples. Six DE comparisons were conducted: 1) the patient versus three healthy male controls; 2) the patient versus three healthy female controls; 3) the mother versus three healthy male controls; 4) the mother versus three healthy female controls; 5) the patient versus the mother; and 6) male controls versus female controls. 11460 expressed genes with a median of over 5 raw CPM (counts per million) for the 8 samples (patient, mother, 3 control males and 3 control females) were included for DE analyses. Absolute expression fold changes of 2 and FDR (false discovery rate) <0.01 were set as the threshold to call genes with significant differential expression (DEGs) (**Table S1-5**). DEGs were used for Gene Ontology (GO) enrichment analysis (http://geneontology.org/). In addition, expression levels of genes in terms of tags per million mapped reads (TPM) and FPKM (fragment per kb exon per million mapped) are shown in **Dataset 2**.

Gene Ontology (GO) overrepresentation test was done by PANTHER (http://geneontology.org/) to test whether upregulated or downregulated DE genes are enriched in GO terms (e.g. biological processes and pathways) compared to the reference of 20996 human genes. FDR<0.05 from Fisher’s Exact test was used for the cutoff. Biological processes or pathways with enrichment fold less than 1 were not shown.

### 2.5 Reverse transcription PCR (RT-PCR)

RT-PCR was done using RNA samples from peripheral blood (SuperScript III Reverse Transcriptase, Invitrogen). Specific cDNA primer pairs are (F: CACTCTTCGCATGGAGTTGA and R: AGTCCACTTGCAGCCACTCT for *F8*, spanning exons 23-25 of NM_000132.3; F: CCAGATGATGGATCAAGGCT and R: CTCTTTTGGGCCTGTATGGA for *BRCC3*, spanning exons 6-7, NM_024332.3).

### 2.6 X-chromosome inactivation (XCI) evaluation

XCI status in the mother was evaluated by the analysis of expressed SNP reads in a few representative X-linked genes that are well expressed and subject to XCI (*ABCD1* and *SLC25A43* in this study; **Table S6**) (Balaton, Cotton, & Brown, 2015). The ratio of SNP read counts of one nucleotide versus total SNP reads at each SNP indicates the chance of this one to be silenced. For a gene subject to XCI, the ratio of SNP read counts of multiple SNPs at one given allele inferred from the SNP information from the patient versus total SNP reads indicates the chance of this allele to be silenced.

To further validate the results, methyl-sensitive PCR-based assay for the *AR* (Androgen receptor) locus containing the polymorphic tri-nucleotide repeat was performed for DNA from mother’s blood or from sister’s saliva sample as previously described with minor modifications (Allen, Zoghbi, Moseley, Rosenblatt, & Belmont, 1992). In brief, DNA samples were PCR amplified for the CAG repeat region (CTG from the minus strand) in the *AR* exon 1 using primers AR-F1: GCTGTGAAGGTTGCTGTTCCTCAT and AR-R1: TCCAGAATCTGTTCCAGAGCGTGC. In parallel, DNA samples were digested with the methyl-sensitive restriction enzyme HpaII followed by PCR. Sanger sequencing with AR-F1 and comparison of the amplified product from DNA with and without digestion were used to determine the presence of polymorphic repeat in test samples and the extent of XCI skewing.

## 3 RESULTS

### 3.1 Clinical features

A 35-year-old man with a history of hypergonadotropic hypogonadism [HH] was referred to the endocrinology clinic for ongoing management of testosterone therapy and was subsequently evaluated in the medical genetics clinic. He reported an initial diagnosis of hypogonadism about 15 years prior, during evaluation for short stature (160 cm, 1^st^ percentile for age; mother’s height 155 cm; father’s height 188 cm). At that time, physical exam demonstrated sparse axillary and pubic hair, small testes and penis, and prepubertal hair distribution. Testosterone replacement was initiated, and his height increased by 5 cm. In the year preceding referral to our clinic, he had discontinued testosterone therapy due to financial constraints. Without testosterone supplementation, he noticed decreased libido, lack of morning erections, low energy and motivation, difficulty concentrating, and difficulty sleeping. Review of systems was positive for depression and anxiety. He had no history of orchitis or testicular trauma, abnormal bleeding, coagulopathy, developmental delay or intellectual disability, and his only previous surgery was carpal tunnel release. He was not taking any medications and did not report any illicit drug use.

He has no family history of hypogonadism, infertility, short stature, early-onset cataracts, or abnormal development. He has a brother (height 173 cm) and sister (height 168 cm), who both went through puberty normally, and his brother has two biological daughters. He has two maternal uncles (both height 178 cm), both of whom have biological children. He is of mixed European ancestry: Scottish, English, Irish, German, and Norwegian.

Laboratory evaluation demonstrated total testosterone 15 ng/dL (250-1150 ng/dL), free testosterone 1.8 pg/dL (35.0-155.0 pg/dL), FSH 81.6 mIU/mL (0.7-10.8 mIU/mL) and LH 42.6 mIU/mL (8.6-61.8 mIU/mL), consistent with HH with >7-fold increase of FSH. Other laboratory results included TSH 1.82 uIU/mL (0.36-3.74 uIU/mL), free T4 0.9 (0.8-1.5 ng/dL), prolactin 4.3 ng/mL (2.5-17.4 ng/mL), IGF-1 by LC/MS 173 ng/mL (53-331 ng/mL), and hematocrit 46.4% (40-52%). Testosterone replacement was re-initiated for treatment of HH. His SNP microarray results (see below) prompted further clinical evaluation. Ophthalmologic exam revealed bilateral cataracts. Magnetic resonance imaging and angiography of the brain showed no evidence of moyamoya disease or other vascular abnormalities, and a transthoracic echocardiogram was normal. Platelet count and activated partial thromboplastin time were normal.

### 3.2 Cytogenetics and Chromosomal SNP Microarray Analysis (CMA) results

G-banded chromosome analysis and karyotyping showed a normal male karyotype (46,XY), ruling out non-mosaic 47,XXY, which is the most common cause of primary HH. Interphase fluorescence in situ hybridization (IFISH) analysis was performed on peripheral blood cells using probes for chromosomes X (DXZ1) and Y (DYZ3). Results showed one signal each for the X and Y chromosomes in all 500 nuclei scored, greatly reducing the probability of mosaicism for XXY. CMA of peripheral blood was performed. Two copy number variants were detected: a minimum 677 kb duplication of chromosome 12q24.33 ([hg38] chr12: 129,318,332-129,994,649), and a minimum 40.5 kb deletion of chromosome Xq28 ([hg38] chrX: 155,028,583-155,069,073). No large regions of copy number-neutral absence of heterozygosity were detected. CMA showed that both the patient’s mother and sister, but neither his brother nor his two maternal uncles, have the same deletion of chromosome Xq28,. This confirms the maternal origin of the deletion in the patient (Figure S1). The 12q24.33 duplication was found only in his sister, suggesting paternal inheritance (Figure S1).

The 677 kb duplication of chromosome 12q24.33 identified in our patient has been reported previously, although it is unclear from CMA results whether it is a direct or inverted tandem repeat, or an insertion elsewhere in the genome with potential for disruption of genes at the insertion site. The duplication contains the first 4 exons of *TMEM132D*, which encodes a transmembrane protein expressed in mature oligodendrocytes. Two similar paternally-inherited duplications have been reported in the Database of Genomic Variation and Phenotype in Humans using Ensemble Resources (DECIPHER), in patients with abnormal emotion/affect or behavior (278910) and moderate intellectual disability (280752). Four similar duplications have been reported in the Database of Genomic Variants (DGV), a database of copy number variants found in control populations. ClinVar lists 3 similar isolated duplications, all classified as of unclear significance, identified from among 29,083 people with developmental delay and intellectual disability (nssv582618, nssv3396370, and nssv579417). Our patient does not have a history of developmental delay or intellectual disability. We consider the 12q24.33 duplication to be an incidental finding and of uncertain significance.

Thus our study focuses on the genetic and expression analyses of the Xq28 deletion in this patient with HH and his family, since Xq28 deletions have been reported in men with growth restriction and HH (Lavin et al., 2016; Miskinyte et al., 2011).

### 3.3 Xq28 deletion breakpoint mapping

Because of gaps between the probes on the microarray, the Xq28 deletion could have included a portion of exon 1 of *F8* (coagulation factor VIII, OMIM# 300841) and/or up to the first 3 exons of *BRCC3* (BRCA1/BRCA2-containing complex subunit 3, OMIM# 300617). To further map the breakpoints of the patient’s deletion, DNA from peripheral blood was amplified by PCR using primers flanking the deletion, followed by Sanger sequencing. The breakpoints were mapped, and the deleted region on Xq28 was 44,806bp (base pairs) in size ([hg18] chrX: 155,025,883-155,070,688) (**Figure 1A**). The proximal breakpoint was located in a highly repetitive sequence SINE element, which is 3159 bp upstream of the transcription start site (TSS) of the main *F8* transcript variant 1 (NM_000132) transcribed from the minus strand. The distal breakpoint was located in the first intron of *CMC4* and *MTCP1* but is 732 bp upstream of the TSS of *BRCC3*. The deleted region therefore contains the entire *FUNDC2* gene and 907 bp upstream of its TSS, and all but the shared exon 1 (5’ UTR) of *CMC4* and *MTCP1*.

**Figure 1.**
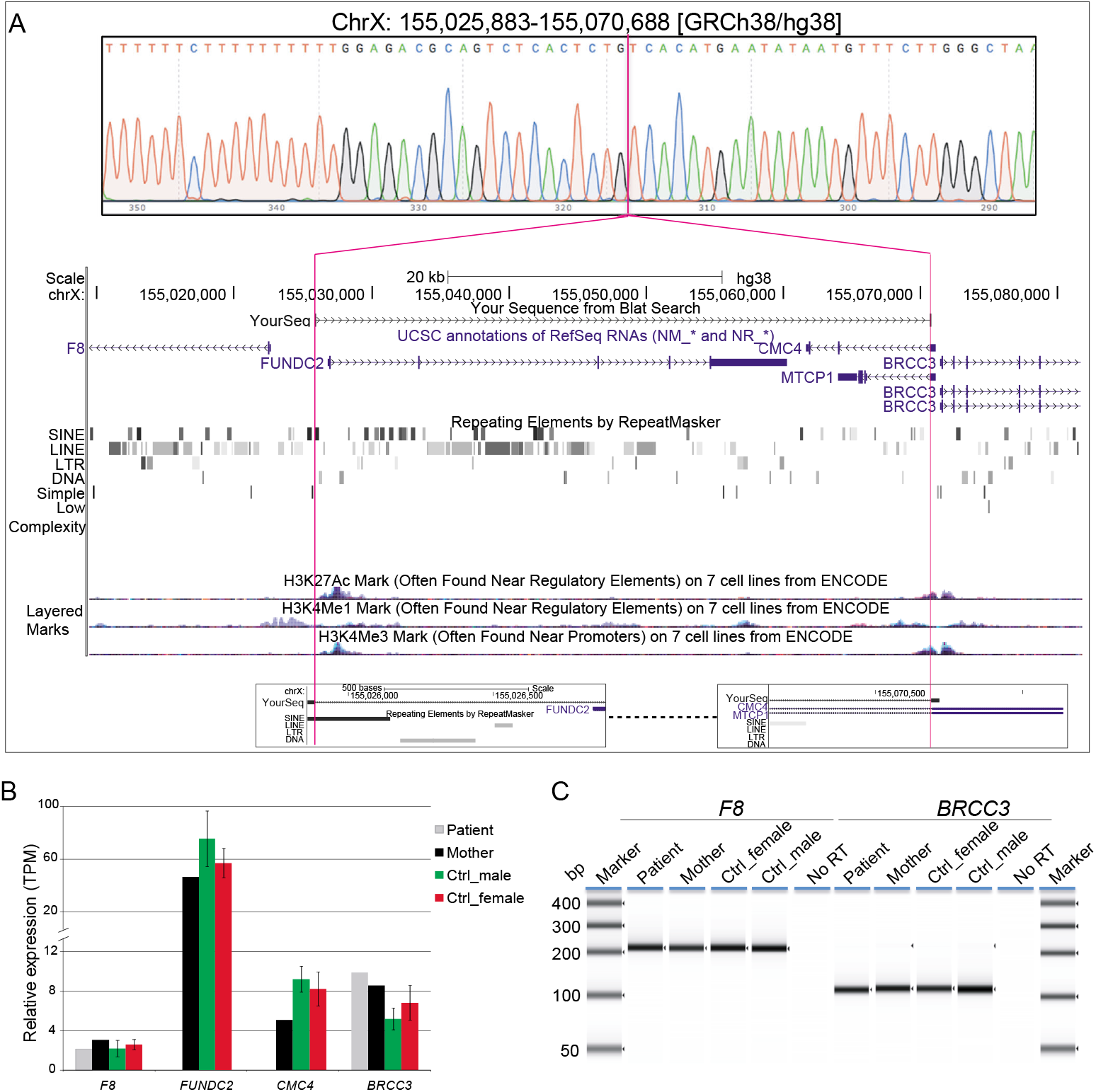
Characterization of Xq28 deletion. (**A**) 44.8 kb Xq28 deletion breakpoints (magenta lines) locate in a SINE element (breakpoint A) and intron 1 of *CMC4* and *MTCP1* (breakpoint B) as well as genes, H3K4me1/3 and H3K27Ac histone marks associated regulatory elements within and near the deletion on Xq28. (**B**) RT-PCR results for the expression of *F8* and *BRCC3* in peripheral blood. (**C**) Relative expression levels of genes within and near the Xq28 deletion measured by RNA-seq. TPM: tags per million. Error bars represent the standard deviation from the mean for three male or female controls.

### 3.4 Effect of the deletion on local gene expression

As the H3K4me1/3 and H3K27Ac histone marks are associated with active regulatory regions such as promoters and enhancers, these elements located in the deleted region could still impact the expression of genes near the deletion (**Figure 1A**). Loss-of-function mutations in *F8* or in *BRCC3* cause hemophilia A or moyamoya angiopathy, respectively (Fujita et al., 2012; Janczar et al., 2014; Lavin et al., 2016; Miskinyte et al., 2011). In addition, BRCC3-containing complexes have been proposed to be involved in regulation of Sertoli cells (Miskinyte et al., 2011). Thus we tested whether expression of *BRCC3* or *F8* was affected in this patient by RT-PCR of peripheral blood. Expression levels of *F8* and *BRCC3* in the patient were similar to his mother and controls (**Figure 1B**).

To further identify dysregulated genes and pathways that could be involved in the HH observed in this patient, we performed transcriptome analysis by mRNA-seq of blood samples from the patient and three of his family members (mother, brother and one maternal uncle). Unfortunately, the quality of RNA-seq data for the brother and the uncle was very poor and thus excluded for DE analysis (see Methods for the detail). By comparing to public mRNA-seq data sets from blood samples of three control males or three control females with age matched to the patient, we confirmed the expression of *FUNDC2* is completely absent in the patient (**Figures 1C** and **S2**; **Table 1**). Since the *CMC4/MTCP1* region has a complex gene structure with a common promoter and 5’ exon which is not deleted in the patient (**Figure 1A**), we carefully examined expression of *CMC4* and *MTCP1*. There is 0 raw RNA-seq count for *CMC4* (Genecode gene ENSG00000182712) after mapping using STAR (**Dataset 1**), which is consistent with the complete lack of *CMC4* gene expression track from the GTEx database of any tissue. Thus *CMC4* is excluded in the DE analysis with EdgeR, which is based on the RNA-seq read counts of Genecode genes (**Table 1**). However, substantial RNA-seq reads were aligned to multiple exons of *CMC4* in RNA-seq of mother and the controls (**Figure S2**), suggesting an annotation problem for the gene ENSG00000182712 for *CMC4*. In contrast, no read was observed on any exon including the undeleted exon 1 in the patient, probably due to degradation of the premature transcript. In support of this, TPM values calculated based on the transcript annotation were obtained for mother and the controls but not the patient (**Figure 1C** and **Dataset 2**). *MTCP1* only has substantial reads on exon 1 that is shared with *CMC4* in mother and the controls (**Figure S2**). The survey of the spliced reads covering the exon-exon conjunction clearly shows a dominant splicing event from exon1 to the next exon of *CMC4* but not *MTCP1* (**Figure S2B**). This strongly suggests that *MTCP1* is not expressed, consistent with the previous findings of no *MTCP1* expression in normal tissues (Madani et al., 1996). We next examined expression of the deletion-flanking genes *BRCC3* and *F8*. No significant expression decrease was observed in the patient versus male or female controls (**Figure 1C** and **Table 1**), confirming the RT-PCR results. Note that expression of BRCC3 is ~2-fold higher in the patient versus control males (FDR=0.03) or females (FDR=0.11) but very similar compared to the mother (Log_2_FC=0.06; **Table S5**).

**Table 1.** Expression changes of genes within or around chromosome Xq28 deletions

| Gene | Patient versus control males |  |  |  | Patient versus control females |  |  |  |
| --- | --- | --- | --- | --- | --- | --- | --- | --- |
|  | log2_FC | log2_CPM | P-Value | FDR | log2_FC | log2_CPM | P-Value | FDR |
| *F8* | -0.42 | 2.43 | 0.44 | 0.67 | -0.13 | 2.23 | 0.84 | 0.95 |
| *FUNDC2* | -12.3 | 4.95 | 6E-27 | 6E-24 | -11.8 | 4.36 | 8E-28 | 2E-24 |
| *BRCC3* | 1.17 | 3.71 | 0.004 | 0.03 | 1.02 | 3.82 | 0.01 | 0.11 |
| Gene | Mother versus control males |  |  |  | mother versus control females |  |  |  |
|  | log2_FC | log2_CPM | P-Value | FDR | log2_FC | log2_CPM | P-Value | FDR |
| *F8* | 0.44 | 2.64 | 0.32 | 0.55 | 0.75 | 2.47 | 0.08 | 0.31 |
| *FUNDC2* | -0.32 | 5.29 | 0.45 | 0.67 | 0.31 | 4.85 | 0.37 | 0.68 |
| *BRCC3* | 1.09 | 3.67 | 0.007 | 0.05 | 0.95 | 3.78 | 0.02 | 0.14 |

Differentiation expression analysis was performed using EdgeR. FC - fold changes, CPM - counts per million.

Taken together, we found that expression of *FUNDC2* and *CMC4* is completely abolished, whereas expression of the deletion-flanking genes *F8* and *BRCC3* were normal or slightly increased in the patient. This is consistent with the absence of hemophilia A or Moyamoya disease in the patient and also suggests a link between loss of function of *FUNDC2* and *CMC4* and the phenotypes of HH observed in the patient.

We further examined whether the deletion has any effect on local gene expression in the mother, who is heterozygous for this deletion. A slight but not significant increase in expression of *BRCC3* and *F8* was observed in the mother compared to control males and females (**Figure 1C** and **Table 1**), consistent with the expression pattern in the patient. Expression of *FUNDC2* in the heterozygous mother is very similar to that in control males and females (log_2_FC=-0.32 and 0.31, respectively; **Table 1**). Interestingly, expression of *CMC4* shown by TPM is 38% and 45% lower than that in female or male controls, respectively (**Figure 1C**). However, these two expression fold changes are not significant as analyzed by Fisher’s exact test based on RNA-seq read counts of *CMC4* transcripts (FDR = 0.8 and 0.5, respectively).

The similar expression of *FUNDC2* between the mother and controls prompted us to examine the XCI pattern. Females inactivate one of their two X chromosomes to achieve a similar expression level as males with one active X (Deng et al., 2014). While random XCI is present in control females, non-random or skewed XCI often occurs in females with an X chromosome with pathogenic deletions.. The extent of XCI skewing is correlated with the pathogenicity of the deletion and also has an effect on expression of mutated affected genes. XCI status was evaluated by expressed SNPs in the mother and the patient by RNA-seq reads (**Table S6**). The X chromosome carrying the Xq28 deletion and inherited by the patient is almost completely skewed to be inactivated (>95%). Independent examination of XCI in blood DNA using the methylation-sensitive enzyme (HpaII) assay of the AR polymorphic repeat locus also shows 91% of XCI skewing toward the deleted X chromosome in the mother (**Figure S3**). Examination of saliva DNA from the patient’s sister, who also inherited the deletion from the mother, shows 82% of XCI toward the deleted X chromosome (**Figure S3**). The skewed XCI in the heterozygous mother and sister strongly suggest the importance of *FUNDC2* and/or *CMC4*. In addition, *FUNDC2* is subject to XCI (Balaton et al., 2015). Expression of the wild-type *FUNDC2* allele that is active in cells would be similar between the mother and control females, consistent with our RNA-seq analysis. However, it is not known whether *CMC4* escapes XCI (Balaton et al., 2015). If *CMC4* escapes XCI in control females, its expression would be lower in the mother compared to control females. Indeed, expression of *CMC4* (5 TPM) is 38% lower than that in control females (7-10 TPM) although this is not significant (FDR = 0.8). Thus XCI skewing occurs in the mother and sister, which is probably adapted at early development to avoid major deleterious effect of this Xq28 microdeletion containing *FUNDC2* and *CMC4*. However, reduction of *CMC4* expression in the mother still could have some effects on gene regulation and development.

### 3.5 Effect of the deletion on global gene expression

To address the effect of loss of function of *FUNDC2* and *CMC4* on global gene expression, we identified a list of differentially expressed genes (DEGs) in the patient compared with control males or females, using a cutoff of FDR<0.01 and absolute fold change >2 (see Methods). A total of 650 and 300 genes are up- or down-regulated respectively in the patient versus control males (**Figure 2A** and **Table S1**). Gene ontology (GO) analysis was performed for up-regulated or down-regulated DEGs separately, which has been shown to be more powerful than using all DEGs together to identify disease-associated pathways (Hong, Zhang, Li, Shen, & Guo, 2013). Indeed, we find that upregulated DEGs are highly enriched in biological processes relevant to the patient phenotype (HH and poor growth), including establishment of Sertoli cell barrier (SCB; also known as blood-testis barrier), apoptosis, cellular response to stimulus and ER stress, inflammatory response, and cell differentiation and proliferation (**Figure 3A** and **Table S7**). In addition DEGs are also relatively enriched (a 2-7 fold enrichment) in mitochondrion function, including positive regulation of mitochondrial membrane permeability, mitochondrial ATP synthesis coupled electron transport, protein localization to mitochondrion, and mitochondrion organization (**Table S7**). Overexpression of genes involved in mitochondrial function could be a way to compensate for the loss of two mitochondrial proteins encoded by *FUNDC2* and *CMC4*, respectively. Strikingly, no enrichment was observed for downregulated DEGs, except for minor enrichment in immunoglobulin production (**Figure 3A**).

**Figure 2.**
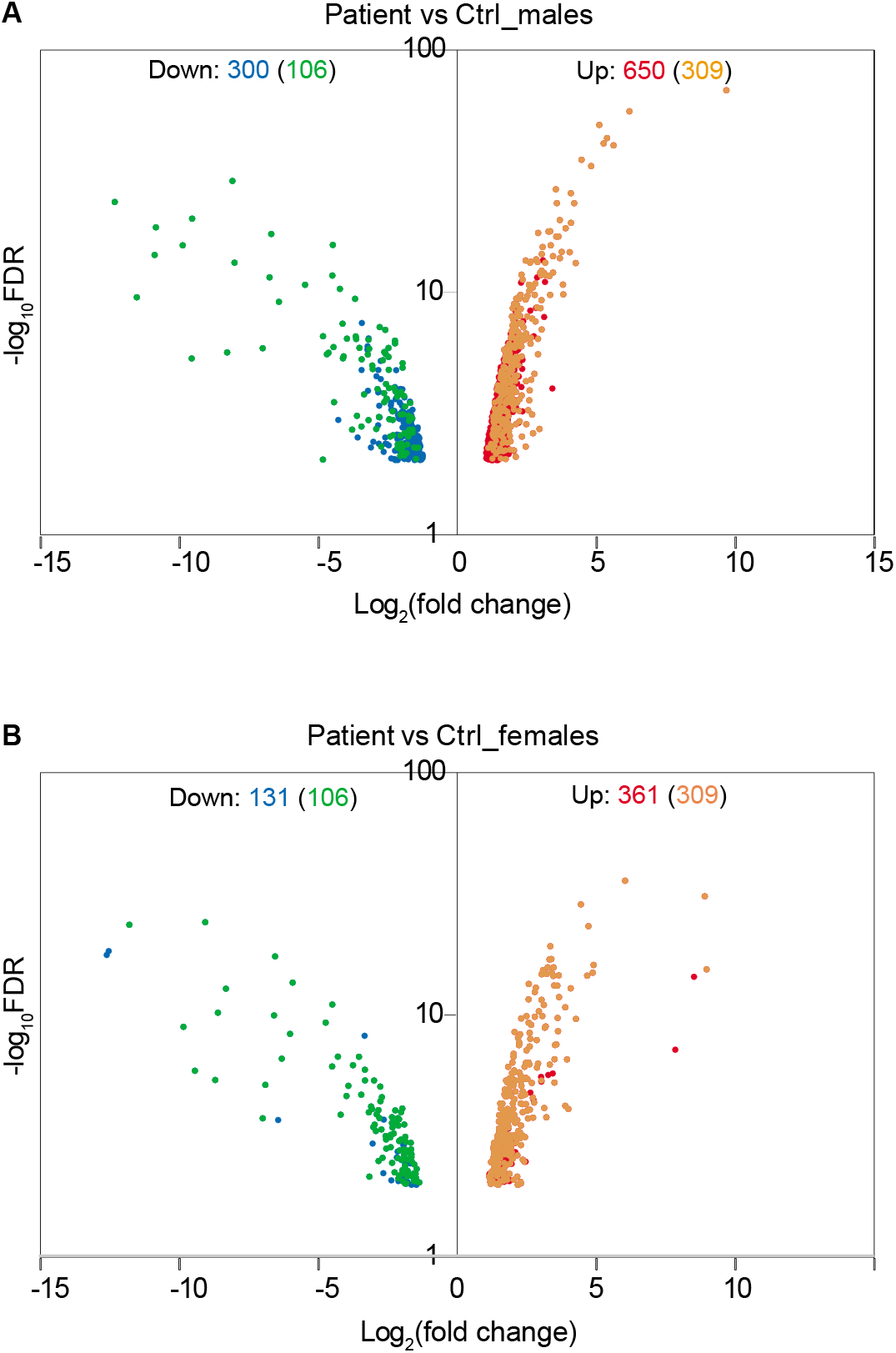
Effect of Xq28 deletion on global gene expression in the patient. (**A**) Volcano diagram of differentially expressed genes (DEGs) in the patient versus control (Ctrl) males. Horizontal axis represents expression fold change changes (log2) and vertical axis represents FDR (log10). DEGs with |log2 fold change| >1 and FDR <0.01 were plotted. Down- or up-regulated DEGs are shown in blue or red, respectively. Overlapped DEGs that were significantly down- or up-regulated both in the patient versus control males and in the patient versus control females are also indicated by green or yellow, respectively. The number of these overlapped DEGs is present in the parenthesis. (**B**) Volcano diagram of DEGs in the patient versus control females. The same analysis is done as in (**A**).

**Figure 3.**
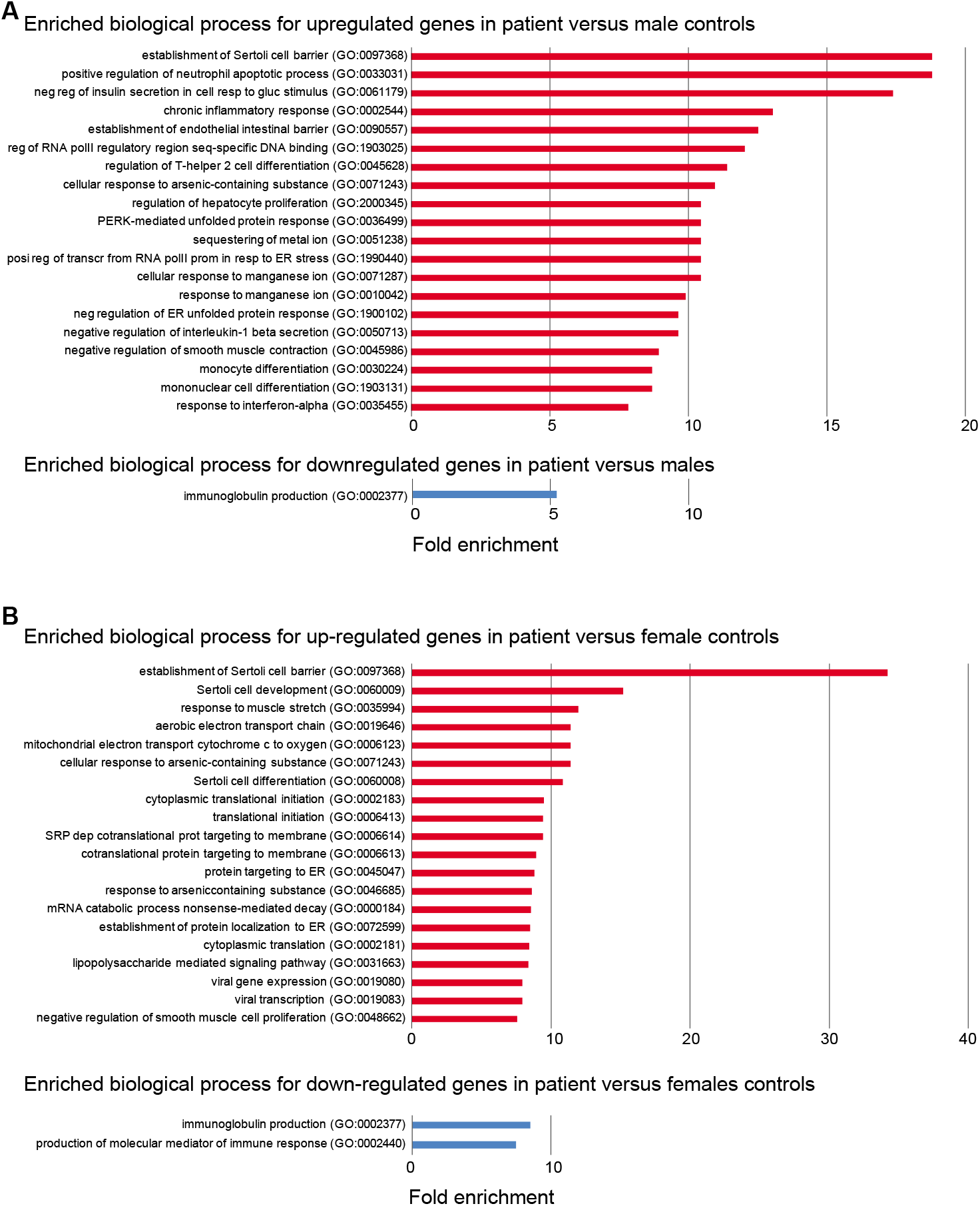
Enriched biological process for up- or down-regulated genes in the patient. (**A**) Enriched biological process from Gene Ontology (GO) analysis for up- or down-regulated in the patient versus control males. The top 20 enriched biological process are shown. FDR<0.05 was used for the cutoff. Biological process with enrichment fold less than 1 were excluded. (**B**) Enriched biological process from GO analysis for down-regulated in the patient versus control females. The same analysis is done as in (**A**).

In the patient versus control females, a total of 361 and 131 genes are up- or down-regulated, respectively (**Figure 2B** and **Table S2**). Interestingly, these DEGs overlapped with those in the patient versus control males (Figure 2B), and the enrichment pattern is also similar (**Figure 3B**). This observation again suggests that up-regulated DEGs in the patient are highly associated with the effect of loss of function of *FUNDC2* and *CMC4* and the phenotype. Intriguingly, three (*ARID4A, ARID4B*, and *ICAM1*) out of five genes involved in regulation of SCB were significantly up-regulated in the patient compared to the controls (**Table S8**). SCB allows Sertoli cells to control the adluminal environment in which germ cells develop by influencing the chemical composition of the luminal fluid. Damage of SCB is associated with spermatogenesis failure and testicular microenvironment deregulation (Lie, Cheng, & Mruk, 2013).

Furthermore, GO pathway analysis of the 650 upregulated genes in the patient versus control males consistently showed that Toll receptor signaling, apoptosis, CCKR (cholecystokinin receptor) signaling - a pathway implicated in digestion, appetite control and body weight regulation - gonadotropin-releasing hormone receptor, and inflammation mediated by chemokine and cytokine signaling pathways are enriched (**Table S9**). Almost all the top ten up-regulated genes (>16-fold up) are involved in one or more of these pathways (**Table S1**). The core up-regulated gene shared by all these pathways is *JUN*, an oncogene that encodes a protein regulating cellular proliferation and preventing apoptosis induced by TNF, which is also up-regulated in the patient. Interestingly, one previous study for testis transcriptome analysis of adult azoospermic patients with 47,XXY (i.e. the most common constitutional cause of HH) and control males by microarray has identified a similar number of DEGs with 656 up-regulated and 247 down-regulated genes in 47,XXY (D’Aurora et al., 2015). This study also reported that up-regulated DEGs in men with 47,XXY are specifically enriched in biological functions and networks, including apoptosis, inflammatory response, hormone regulation and steroidogenesis, and regulation of SCB. The overlapped pathways affected in testis of men with 47,XXY and in the blood of our HH patient with the Xq28 deletion strongly support the idea that disruption of such a gene network involved in apoptosis, inflammation and SCB regulation will lead to HH. However, the genetic cause of HH is very different between patients with 47,XXY and our patient, who has loss of function of *FUNDC2* and *CMC4*. Indeed, only 42/650 (6.5%) and 5/300 (1.7%) of DEGs in our patient are also up- or down-regulated in patients with 47,XXY, respectively (**Table S10**). One central node gene in the top network for the up-regulated DEGs in 47,XXY is *SMAD3*, which is not affected in our patient (**Table S1**). In addition, four DEGs (*ARID4A*, *ARID4B*, *ICAM1* and ATRX) involved in regulation of SCB and SC development and differentiation and identified in our patient are not up-regulated in patients with 47,XXY. Interestingly, *JUN*, the core gene shared by the disrupted pathways in our patient (**Table S9**), is a node gene in the top network for the up-regulated DEGs in patients with 47,XXY. Consistently, four up-regulated genes in the same pathways with *JUN* in the patient (*DUSP1, MAP4K4, TNFAIP3* and *TUBAIA*) are also up-regulated in patients with 47,XXY, suggesting a partial overlap of targeted genes in the etiology of HH.

We also performed DE and GO analyses for the heterozygous mother in whom the deleted X chromosome is skewed to be almost completely inactivated. Interestingly, a similar set of DEGs was identified in the mother versus controls (**Figures S4A-B** and **S5**, and **Tables S3** and **S4**), suggesting that deletion of *FUNDC2* and/or *CMC4* still has an effect on gene regulation in the mother. In contrast, only 16 DEGs were identified between control males and females (**Figure S4C**). While *FUNDC2* is subject to XCI, *CMC4* could escape XCI and contribute to its expression levels in control females. In addition, a small portion of cells in the mother still could keep the deleted X chromosome active since the skewing extent of XCI is ~91%. Indeed, a relative lower expression of *CMC4* (38% down) is observed in the heterozygous mother compared to control females, which could be related to such gene dysregulation. In support of this, DE comparison of the patient and his mother only showed a small number of genes (10 up and 53 down) with expression changes (**Table S5**).

Overall, these findings are relevant to the phenotype of our patient with HH in whom FSH but not LH is highly elevated, and suggest a role of *FUNDC2* and *CMC4* in regulation of apoptosis and inflammation.

## 4. DISCUSSION

We describe a 44.8 kb deletion of chromosome Xq28 encompassing the entirety of *FUNDC2*, exons 2-3 of *CMC4*, and exons 2-5 of *MTCP1*. This deletion has not previously been reported in the DGV, DECIPHER, or ClinVar. However, Xq28 deletions involving similar regions have previously been described in the medical literature, and associated phenotypically with either hypergonadotropic hypogonadism [HH] and Moyamoya disease, or severe hemophilia A and Moyamoya [SHAM] syndrome (**Tables 2** and **S11**). Miskintye et al. described findings in 7 patients from 2 families with syndromic Moyamoya disease with HH, and a third family with syndromic Moyamoya disease with no information on HH (**Table 2**) (Miskinyte et al., 2011). The critical region of overlap for syndromic Moyamoya disease among these 3 families encompasses exon 1 of *CMC4* and *MTCP1* and the first 3 exons of *BRCC3* (**Table S11**). *BRCC3* encodes a deubiquitinating enzyme as a member of two *BRCA1* and *BRISC* complexes. Further experiments in this study using zebrafish models demonstrated that deletion of *BRCC3* but not *CMC4* results in defective angiogenesis, further confirming the *BRCC3* role in Moyamoya disease. In addition, *BRCC3* was proposed by Miskintye et al. to play a role in HH, since *BRCC45*, a member of the BRCC3-containing complexes, is highly expressed in germ cells and SCs.

**Table 2.** Clinical features of patients reported with chromosome Xq28 deletions

|  | Pt<br>(this<br>study) | Lavin et<br>al. (2016) | Janczar<br>et al.<br>(2014) | Fujita<br>et al.<br>(2012) |  | Miskinyte et al. (2011) |  |  |
| --- | --- | --- | --- | --- | --- | --- | --- | --- |
|  |  |  |  | Pt<br>1 | Pt<br>2 | Family 1 | Family<br>2 | Family 3 |
| Age (years) | 35 | 37 | 10 | 17 | 25 | 28-48 | 14,17 | 18, 26 |
| Short stature | + | + | NR | + | - | 5 of 5 | 2 of 2 | 2 of 2 |
| Premature<br>gray hair | - | + | NR | NR | NR | 5 of 5 | 0 of 2 | 1 of 2 |
| Eye findings | Bilateral<br>cataracts | Unilateral<br>proptosis | NR | NR | NR | Bilateral<br>early-onset<br>cataracts<br>(4 of 5) | 0 of 2 | NR |
| Facial<br>dysmorphism | - | NR | + | NR | NR | 5 of 5 | 2 of 2 | 1 of 2 |
| Moyamoya<br>disease | - | -<br>(PCA<br>aneurysm) | + | NR | NR | 4 of 5 | 2 of 2 | 2 of 2 |
| Cardiac | Normal | NR | NR | NR | NR | DCM<br>(3 of 5),<br>Isolated<br>LVE (2<br>of 5) | 0 of 2 | MI,<br>Cardiomegaly<br>(1 of 2) |
| Hemophilia<br>A | - | + | + | + | + | 0 of 5 | 0 of 2 | 0 of 2 |
| HH | + | + | Unknown | NR | NR | 5 of 5 | 2 of 2 | NR |

Pt – patient, HH - hypergonadotropic hypogonadism DCM - dilated cardiomyopathy, LVE – left ventricular enlargement, MI – myocardial infarction, NR - Not reported, PCA - posterior communicating artery, TIA – transient ischemic attacks

Loss-of-function variants in *F8*, which encodes coagulation factor VIII, cause hemophilia A, and microdeletions at Xq28 have been identified in patients with SHAM syndrome. Lavin et al. described a 37-year-old man with hemophilia A, who presented with left knee arthropathy and was noted to have short stature, sparse body hair, gynecomastia, prematurely gray hair, and unilateral proptosis (Lavin et al., 2016). He was also found to have HH and a posterior communicating artery aneurysm on brain imaging. He was described as having SHAM syndrome (**Table 2**). Genetic evaluation revealed a 201 kb deletion of Xq28 that removed exons 1-14 of *F8* and the entirety of *FUNDC2, CMC4, MTCP1*, and *BRCC3* (**Table S11**). Janczar et al. described SHAM syndrome in a 10-year-old boy with hemophilia A who presented with focal neurologic deficits and was found on MRI to have ischemic stroke and Moyamoya disease (**Table 2**) (Janczar et al., 2014). Genetic evaluation revealed a 150 kb deletion of Xq28 involving exons 1-6 of *F8* and the entirety of *FUNDC2, CMC4, MTCP1*, and *BRCC3* (**Table S11**). Fujita et al. reported three patients with severe hemophilia with atypical intron 22 inversions and large deletions of *F8*. In 2 of the patients, the deletion encompassed genes other than *F8* (**Table S11**) (Fujita et al., 2012). Evaluations for Moyamoya disease or HH were not reported in either patient. Genetic evaluation revealed a microdeletion involving exons 1-22 of *F8* and the entirety of *FUNDC2*, *CMC4*, and *MTCP1* in both patients. The deletion extended past *BRCC3* in one of them (**Table S11**).

Of the clinical features described in these previous reports of Xq28 deletions, our patient has HH, short stature, and bilateral cataracts on ophthalmologic exam. He has no evidence of coagulopathy or hemophilia A, cardiomyopathy, or Moyamoya disease on further lab tests and imaging. This phenotype is consistent with expression analysis, since *BRCC3* and *F8* expression were normal in the peripheral blood of the patient. The critical minimal region of overlap between our patient and the previously reported 8 patients is now narrowed to exonic deletions of *CMC4* and *MTCP1*. Since *MTCP1* is not expressed in normal tissues, it is possible that *CMC4* plays a role in HH. Our expression analysis showed that up-regulated genes in the patient are enriched in SC regulation, gonadotropin-releasing pathway, apoptosis, and inflammatory response. This is strongly consistent with the extremely high level of FSH in the patient. It is not clear how loss of function of *CMC4* causes this, due to very limited functional studies of this gene. One recent study using the nude mouse tumor model shows that suppression of *Cmc4* (misnamed as *Mtcp1* in this study, since the antibody used is for the peptide encoded by *Cmc4*) by *miR-126* is involved in repression of tumor growth and migration (Han, Liu, Wu, & Liu, 2018). This suggests a role of *CMC4* in cell proliferation and mobility. Given that the CMC4 protein level is highest in fetal testis among the examined tissues including adult testis in HIPED, it is tempting to speculate that loss of function of *CMC4* contributes to HH in men. How *CMC4* is associated with genes involved in SC regulation, gonadotropin-releasing pathway, apoptosis, and inflammatory response is not clear.

Although the minimal overlapped region between our patient with HH and those from the literature doesn’t include *FUNDC2*, we cannot exclude a possible role of *FUNDC2* in HH, since those deletions also contain *BRCC3* that could be involved in HH. *FUNDC2* is also a mitochondria protein, like *CMC4*, and regulates platelet lifespan by preventing apoptosis under hypoxia stress (Ma, Zhu et al., 2019). An important paralog of *FUNDC2* is *FUNDC1*, which is highly conserved and regulates mitophagy and inflammatory response. It has been recently shown that the ortholog of *FUNDC1* in C. elegans (no *FUNDC2* in C. elegans) is strongly expressed in sperm and contributes to paternal mitochondria elimination in sperm (Lim et al., 2019). The most likely interacting protein for FUNDC2 is FUNDC1 as predicted by the STRING Interaction Network in the GeneCards database, suggesting functionary conservation. Thus FUNDC2 in humans could be also involved in mitophagy, apoptosis/inflammation, and spermatogenesis.

Our patient also has bilateral cataracts. Eye defects are observed in the previously reported patients with a deletion of *FUNDC2*, but not with deletion of *CMC4* and *BRCC3* (**Table 2** and **S11**), suggesting the involvement of *FUNDC2* in eye disease. Interestingly, we find that 5 out of 9 genes (*ENY2, HMGCR, PIM3, MAP4K4* and *KLF4*) involved in negative regulation of insulin secretion in cellular response to glucose stimulus (GO:0061179; **Figure 3**) are upregulated in the patient. This suggests a possible link of dysregulation of insulin pathways and cataracts in the patient, since cataracts are one of the sight-related complications of diabetes. In support of this, three genes (*KLF4, RIT1*, and *VEGFA*) involved in post-embryonic camera-type eye development (GO:0031077 that is not included in the enriched biological process) are also significantly upregulated in the patient (**Table S1**). These genes are all related with cell growth/apoptosis regulation.

## 5 CONCLUSION

We describe a rare case of syndromic hypergonadotropic hypogonadism [HH] with short stature and cataracts in a 35-year-old man, in whom we identified a 44.8 kb microdeletion of chromosome Xq28 involving the entirety of *FUNDC2* and most of *CMC4* and *MTCP1*. Further genetic and expression analyses suggest a novel link between *CMC4* and/or *FUNDC2* and apoptosis, inflammation and FSH and SCB regulation in men, which is strongly supported by the finding that the disrupted pathways of upregulated DEGs overlap between our patient and men with 47,XXY. In addition, *CMC4* and/or *FUNDC2* are probably important in female-specific development, since both female family members (the mother and sister) who carry this deletion and have no clinical abnormalities have a very high level of XCI skewing. Taken together, our study provides novel insights into how loss of *CMC4* and/or *FUNDC2* contributes to dysregulation of apoptosis, inflammatory response, and HH and the avenue for studies of molecular mechanisms of HH.

## ACKNOWLEDGENTS

We thank the patient and his family, without whom this work would not have been possible. We thank Drs. Yu Wu and Wenjing Wang for their technical support. This work is partly supported by National Institutes of Health (NIH)/National Institute of General Medical Sciences (NIGMS) Grant R01GM127327-01 (to X. Deng).

## Data Availability Statement

The data that support the findings of this study are available on request from the corresponding author. The data are not publicly available due to privacy or ethical restrictions.

## Supporting information

**Figure S1. Family pedigree** Circles indicate females, and squares indicate males. Diagonal lines designate deceased family members. Arrow indicates the proband. Remaining details are defined in the Key..

**Figure S2. RNA-seq reads mapped at the Xq28 deletion region show that expression of *FUNDC2* and *CMC4* is completely abolished in the patient**

(**A**) Mapped RNA-seq reads were binned in 100bp windows and visualized by the Integrative Genomics Viewer (IGV). Accumulated reads are exclusively located at exons of *FUNDC2, CMC4* and *BRCC3* in the heterozygous mother and controls. Only the first exon of *MTCP1* shared with *CMC4* has substantial reads aligned. In the patient, there is a complete loss of reads at *FUNDC2* and *CMC4* but not *BRCC3*. (**B**) Individual reads visualized by IGV. Spliced reads spanning two exons were almost mapped to the exon-exon junctions for *CMC4* in the mother, suggesting that *CMC4* but not *MTCP1* is expressed. No reads are observed at *CMC4* in the patient, consistent with the loss of this gene.

**Figure S3. Skewed XCI patterns in the heterozygous mother and sister are revealed by the methyl-sensitive PCR-based assay**

(**A**) PCR results for the AR (Androgen receptor) locus containing the polymorphic tri-nucleotide repeat (CTG from the minus strand as shown here). Genomic DNA from mother’s blood or from sister’s saliva sample before and after digestion with the methyl-sensitive restriction enzyme HpaII was used for PCR. (**B**) Snap shots of Sanger sequencing results for the PCR products of the AR repeat regions in the mother. Before HpaII digestion, the sample from the mother starts to show double peaks (T/A) at the position where the two alleles have different length of CTGs. This information is used to infer the number of repeats ((CTG)n) for each allele, which shows (CTG)19 : (CTG)22 for the mother. After HpaII digestion, the allele on the inactive X which is methylated can’t be digested and becomes PCR amplified and sequenced. The ratios of the peak heights between T and A at the 1^st^ and 2^nd^ positions of double peaks before and after digestion is compared to measure the ratio of XCI for each allele (e.g. similar ratios mean equal chance of XCI and increased/decreased ratios, skewed XCI). In the mother, these ratios are decreased by 10-fold after digestion, suggesting that the X chromosome with longer repeats is preferred for digested (i.e. the other X with (CTG)19 is preferred for XCI). (**C**) Snap shots of Sanger sequencing results for the PCR products of the AR repeat regions in the sister. Similar analysis was done as in (**A**). This shows (CTG)19 : (CTG)24 for the sister, suggesting the X chromosome with (CTG)19 is inherited from the mother and has the Xq28 deletion. Consistently, this deleted X chromosome is also preferred for XCI in the sister.

**Figure S4. Effect of Xq28 deletion on global gene expression in the mother**

(**A**) Volcano diagram of differentially expressed genes (DEGs) in the mother versus control (Ctrl) males. Horizontal axis represents expression fold change changes (log2) and vertical axis represents FDR (log10). DEGs with ļlog2 fold changeļ >1 and FDR <0.01 were plotted. Down- or up-regulated DEGs are shown in blue or red, respectively. Overlapped DEGs that were significantly down- or up-regulated both in the mother versus control males and in the mother versus control females are also indicated by green or yellow, respectively. The number of these overlapped DEGs is present in the parenthesis. (**B**) Volcano diagram of DEGs in the mother versus control females. The same analysis is done as in (**A**). (**C**) Volcano diagram of DEGs in control males versus control females. The same analysis is done as in (**A**).

**Figure S5. Enriched biological process for up- or down-regulated genes in the mother**

(**A**) Enriched biological process from Gene Ontology (GO) analysis for up- or down-regulated in the mother versus control males. The top 20 enriched biological process are shown. FDR<0.05 was used for the cutoff. Biological process with enrichment fold less than 1 were excluded. No any enrichment was observed for down-regulated genes. (**B**) Enriched biological process from GO analysis for down-regulated in the patient versus control females. The same analysis is done as in (**A**).

**Table S1. Differential expression changes between the patient and control males**

**Table S2. Differential expression changes between the patient and control females**

**Table S3. Differential expression changes between the mother and control males**

**Table S4. Differential expression changes between the mother and control females**

**Table S5. Differential expression changes between the patient and mother**

**Table S6. Skewed XCI in the mother shown by analysis of expressed SNPs by RNA-seq in blood**

**Table S7. Enriched biological process for upregulated DE genes in the patient versus control males**

**Table S8. Four upregulated DE genes in the patient are involved in sertoli cell regulation in GO biological process analysis**

**Table S9. Enriched pathways for upregulated DE genes in the patient versus control males**

**Table S10. A list of genes up- or down-regulated genes in the pateint versus control males and in KS patients versus control males**

**Table S11. Molecular characteristics of Xq28 deletions associated with syndromic forms of hypergonadotropic hypogonadism in men**

**Dataset 1. Raw read counts for 8 RNA-seq samples**

**Dataset 2. FPKM and TPM values for 8 RNA-seq samples**

## Web Resources

Database of Genomic Variants - http://www.dgv.tcag.ca

ClinVar - https://www.ncbi.nlm.nih.gov/clinvar/

DECIPHER - http://decipher.sanger.ac.uk

Gene Tissue Expression Project - http://www.gtexportal.org

Online Mendelian Inheritance in Man - http://www.omim.org

UCSC Genome Browser - http://genome.ucsc.edu

Bioconductor - http://www.bioconductor.org/

